# Geographic variation in the songs of two closely related song-learning species, Anna’s and Costa’s hummingbird (*Calypte anna, C. costae*)

**DOI:** 10.1101/2023.11.26.568754

**Authors:** A. N. Berger, W. Y. Ye, A. S. Padilla, B. C. Gumbi, C. J. Clark, P. Campbell

## Abstract

In species that learn their song, cultural transmission of song components can lead to the accumulation of variants that differ among populations, resulting in the formation of dialects. Three avian clades are thought to have independently evolved song learning – parrots, oscine passerines, and hummingbirds. Dialects have mainly been studied in passerines. We extend the study of dialects to the bee hummingbird clade, focusing on Anna’s and Costa’s hummingbirds (*Calypte anna* and *C. costae*). Both species are vocal learners. Anna’s produces complex, three phrase, multi-syllable songs and Costa’s produces simple, one phrase songs. We recorded 5-24 males per population (5 Costa’s and 6 Anna’s populations) across the species’ ranges in the Western United States and tested for evidence of geographic variation in song. We found minor population differences in frequency measures of Costa’s song, but song form was invariant across populations. Anna’s song was contrastingly variable with population differences in both syllable use and multiple spectral and temporal measures. The most strongly differentiated Anna’s population in our study, Seattle (Washington State), is the product of a recent northward range expansion facilitated by human activities that provide additional food sources for hummingbirds. The loss and modification of syllables in this population is suggestive of a founder effect on song. This study provides insight into song evolution in non-passerine vocal learners and contributes to understanding of how complex signals evolve.

---

Acoustic communication has intrigued scientists for millennia. (Pliny the Elder, 50 A.D.; Darwin, 1871). Of particular interest are the evolution and mechanistic basis of vocal learning–a defining feature of humans that has evolved independently in a subset of other mammalian taxa (pinnipeds, cetaceans, elephants, bats), and three bird lineages (songbirds, parrots, hummingbirds) (Nottenbohm, 1972; Janik and Slater, 1997; Tyack, 2019; Janik and Knörnschild, 2021). Vocal learning is a complex and imitative social process that requires the capacity to memorize acoustic input and to match acoustic output to this internal template. At every step of the vocal learning and production process there is a potential for errors that generate vocal novelties. Like genetic mutations, these vocal novelties can rise to high frequency and ultimately become fixed in a population, resulting in among population acoustic differences or dialects. (Marler and Tamura, 1964; Lemon, 1975; Baptista, 1977; Marler and Peters, 1987; Slater, 1989).

The study of dialects in oscine birds has a long history. Marler and Tamura set the stage with their foundational study on the white-crowned sparrow (*Zonotrichia leucophrys*), in which they documented stereotyped local dialects in three populations in California (Marler and Tamura, 1962). This work catalyzed an explosion of studies looking at incidences of dialects (Lemon, 1966; Nottebohm, 1969; Baptista and King, 1980; Wang et al., 2022), the adaptive function of dialects (Baker, 1975; Mundinger, 1982; Baker and Cunningham, 1985), and the ecological predictors and mechanisms of dialect formation (Lemon, 1975; Morton, 1975; Slater, 1989; Kroodsma and Miller, 1996; Podos and Warren, 2007; Derryberry, 2009). In parallel, dialect research has expanded to include the two non-oscine lineages of song learners, parrots (Bond and Diamond, 2005; Wright et al., 2005; Kroodsma et al., 2013, reviewed in Wright and Dahlin, 2018) and hummingbirds (Snow, 1968; Wiley, 1971; Baptista and Schuchmann, 1990; Gaunt et al., 1994; González and Ornelas, 2009; Araya-Salas and Wright, 2013. Lara et al., 2015). Lek mating systems are common in hummingbirds and most prior research on hummingbird dialects has focused on vocal variation between leks in species that have well defined lek boundaries. Even at this microgeographic scale, there is evidence for dialects. For example, the song of wedge-tailed Sabrewings (*Campylopterus curvipennis*) has lek specific introductory syllables (González and Ornelas, 2005; González and Ornelas, 2014), and song neighborhoods in long billed hermits (*Phaethornis longirostris*) are grouped by leks (Araya-Salas et al., 2019).

In this paper we move the study of hummingbird dialects to a macrogeographic scale to test for evidence of geographic variation in the song of two hummingbird species, Anna’s and Costa’s hummingbirds (*Calypte anna* and *C. costae*, respectively; hereafter, Anna’s and Costa’s). Anna’s and Costa’s are sister species and the sole members of the *Calypte* genus within the bee (*Mellisuginii*) hummingbird clade (McGuire et al., 2007). Both species learn their songs (Baptista and Schumman, 1990; Johnson and Clark, 2020) and sing during territorial and courtship displays (Wells et al., 1978; Stiles, 1982). Despite being sister taxa, Anna’s and Costa’s have remarkably different song structures. Anna’s song is spectrally complex and multi-syllabic with multiple phrases whereas Costa’s song is pure tone, mono-syllabic, and comprises a single phrase (Fig. 1a and b).

**Figure 1.**
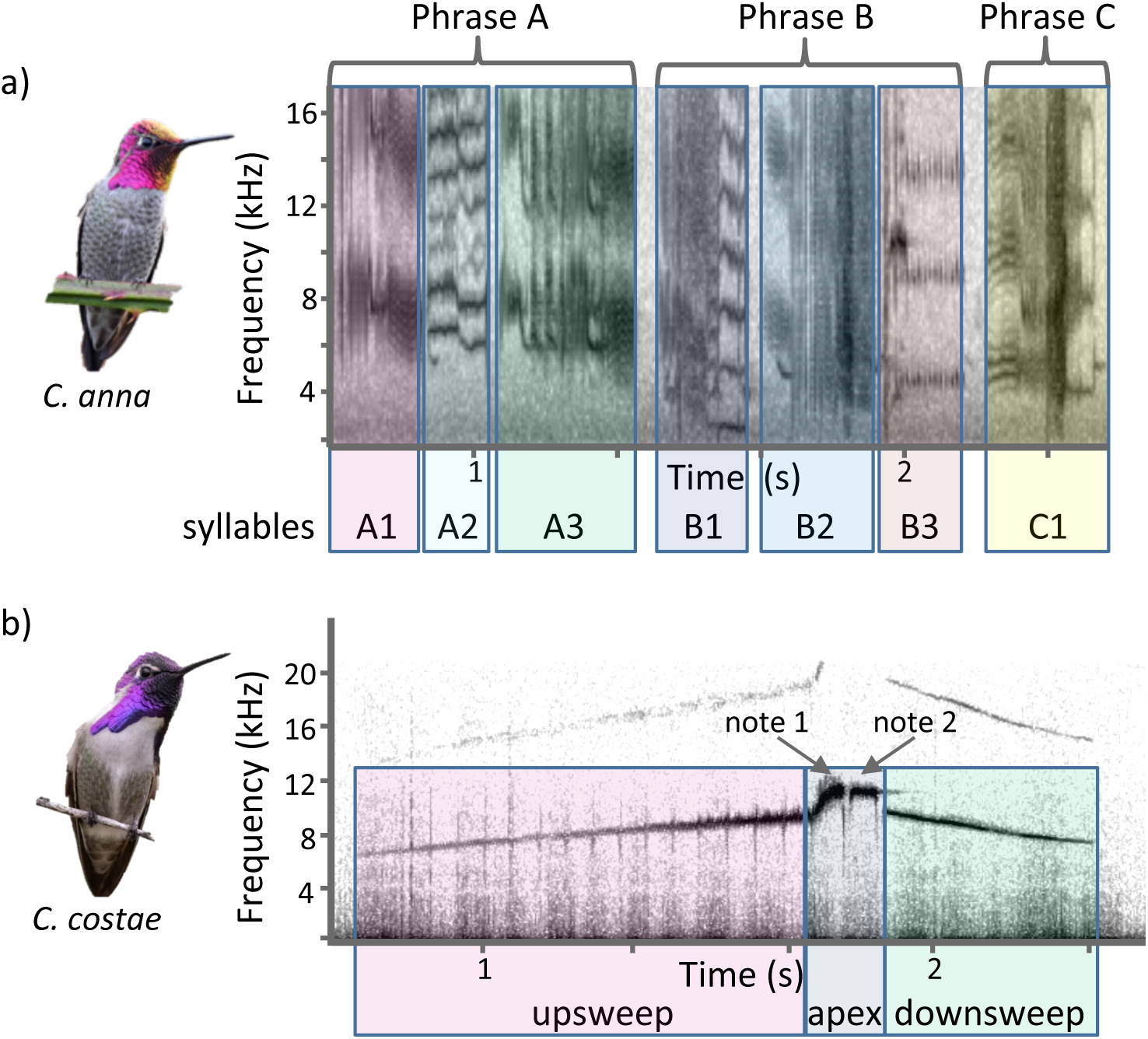
Anna’s and Costa’s song. **a)** Anna’s hummingbirds sing a complex three phrase (A-C) song with multiple syllables (A1-A3, B1-B3, C1). **b)** Costa’s hummingbirds sing a simple one phrase song with three elements: upsweep, two note apex, and downsweep. Images, Steven Mlodinow, Tom Friedel.

As for other hummingbird species, there is evidence for within-population variation in both Anna’s and Costa’s songs (Williams and Houtman, 2008; Yang et al., 2007). The only prior study conducted at a larger spatial scale found acoustic differences between the songs of an isolated Anna’s population from Guadalupe Island (240 km from the nearest mainland) and birds from a mainland population (Mirsky, 1976). However, whether song dialects exist between mainland populations that are not geographically disjunct from one another is unknown.

We recorded Anna’s and Costa’s hummingbirds across their ranges in the Western United States (Fig. 2), and collected spectral, temporal, and qualitative measurements of the songs to characterize the presence or absence of dialects. Given that the complex multi-syllabic nature of Anna’s song provides ample opportunity for improvisation and error in the song learning process, we expected to find clear evidence for dialects in this species. In contrast, the simpler structure of Costa’s song leaves less room for error. Therefore, we expected to detect less, if any, among-population differences in Costa’s song.

**Figure 2.**
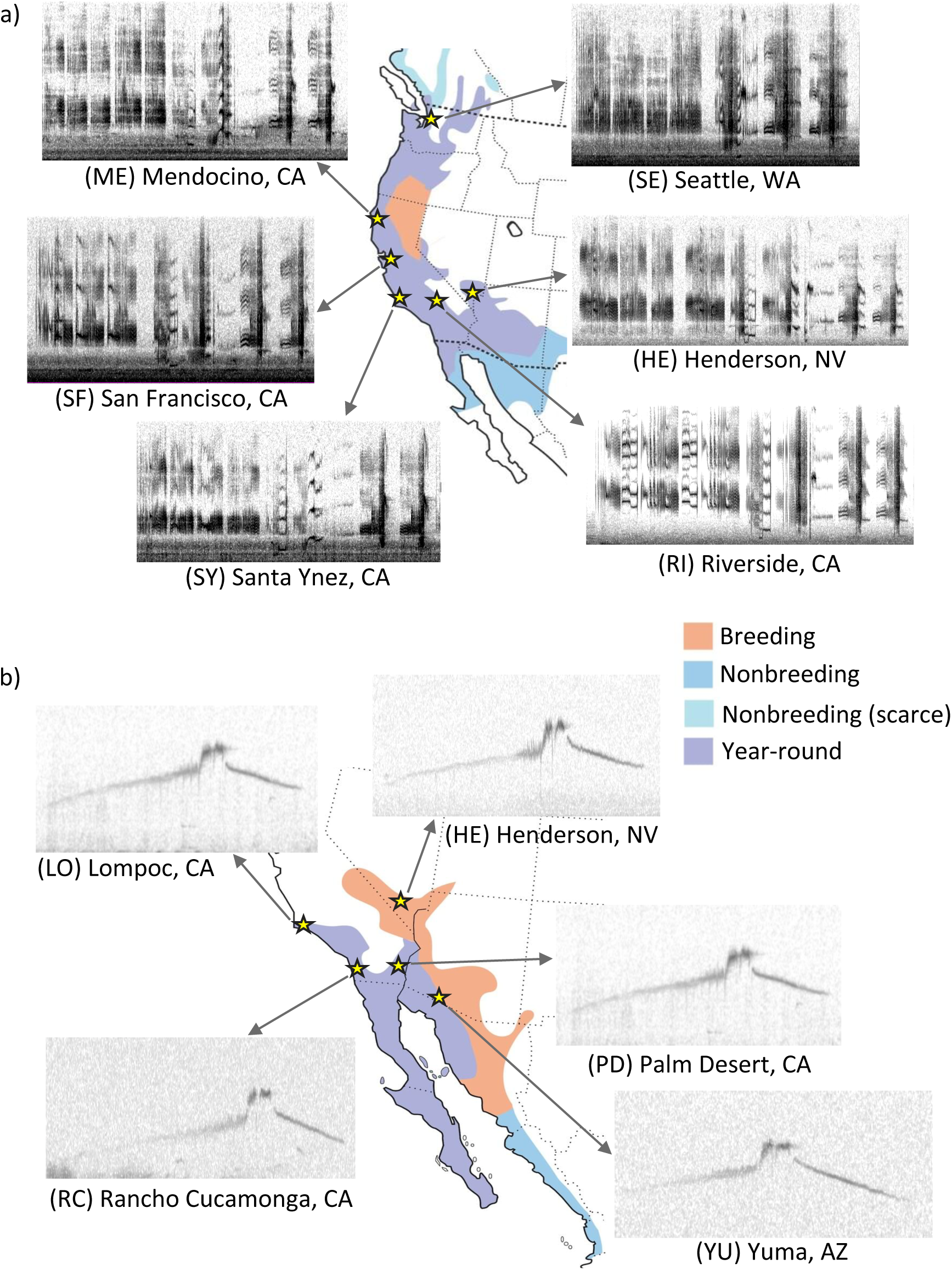
Anna’s and Costa’s geographic distributions and approximate locations of sampling sites with representative spectrograms. **a)** Anna’s sample sizes: RI (Riverside, CA) *n*=15; SY (Santa Ynez, CA) *n*=10, SF (San Francisco, CA) *n*=25; ME (Mendocino, CA) *n*=10; SE (Seattle, WA) *n*=25; HE (Henderson, NV) *n*=5**. b)** Costa’s sample sizes: PD (Palm Desert, CA) *n*=15; RC (Rancho Cucamonga, CA), *n*=7; LO (Lompoc, CA) *n*=10; HE (Henderson, NV) *n*=6. Range maps modified from All About Birds (allaboutbirds.org).

## METHODS

We sampled Anna’s and Costa’s hummingbird song from geographically distinct populations in the Western United States during the January to June breeding season in 2020 and 2021 (Anna’s, *n*=6 populations, 10-20 birds/population; Costa’s, *n*=5 populations, 5-20 birds/population; Fig. 2a and b). The ranges of straight line distances between sampled populations were 171-1,571 km for Anna’s and 116-619 km for Costa’s. We recorded songs from males singing on their breeding territories (5-30 songs/male). Breeding males of both species are territorial and can be identified by perch fidelity (Stiles, 1982). Territories are stable from day to day (Stiles, 1973; Clark and Russell, 2020). Recordings from a given male were collected consecutively over the course of one to two days, with most recordings taken in a single day. Recordings were captured using a Zoom F8 multitrack field recorder (Zoom Corporation, Tokyo, Japan), at a sample rate of 48kHz with a 24-bit depth and a Sennheiser K6 microphone (Sennheiser electronic GmbH & Co. KG, Wedemark, Germany; frequency range 30Hz −20 kHz, ± 1dB frequency handling) with a parabola shell (Wildtronics, LLC Mono Parabolic microphone, Newton Falls, OH, USA) at a distance of ≤15 meters from the focal bird. We divided the songs into phrases, syllables, and elements by visual inspection of the spectrograms generated in Raven Pro v1.6 (Cornell Lab of Ornithology, Cornell University, Ithaca, NY, USA) using a 512 FFT smooth Hamming window.

### Spectral Analysis

#### Anna’s Hummingbird Song

Anna’s song consists of three multi-syllabic phrases (A-C; Fig. 1a). Phrase A consists of 2-4 syllables, phrase B consists of 2-3 syllables, and phrase C consists of 1-2 syllables. The song is highly complex with high entropy and a frequency range of 1.5 – 20 kHz, with most of the energy between 1.5 and 4 kHz. We defined syllables as repeatable components that have an inter-syllable duration less than 0.08 seconds and defined phrases as repeated groupings of syllables with between phrase durations greater than 0.08 seconds. Song syntax is stereotyped within individuals (Yang et al., 2007, ANB personal observation). For each male we chose one representative song with the highest signal to noise ratio and the lowest background noise. For each syllable we measured duration and peak frequency in Raven (Table 1).

**Table 1.** Analysis of variance results for spectral and temporal measures of Anna’s hummingbird song.

|  | Measurement | *n* | *F* | *p* |
| --- | --- | --- | --- | --- |
| **Phrase A** |  |  |  |  |
| Syllables | **A1 Duration (seconds)** | 62 | **3.91** | **0.004** |
| A1-A2 | A1 Peak frequency (kHz) | 62 | 1.54 | 0.190 |
|  | **A2 Duration (seconds)** | 61 | **2.43** | **0.047** |
|  | A2 Peak frequency (kHz) | 61 | 1.88 | 0.113 |
| **Phrase B** |  |  |  |  |
| B1 | **B1 Peak frequency (kHz)** | 47 | **2.94** | **0.030** |
|  | B1 Duration (seconds) | 47 | 2.33 | 0.071 |
|  | Upper whistle (kHz) | 47 | 0.62 | 0.653 |
| B2 | B2 Peak frequency (kHz) | 63 | 2.15 | 0.072 |
|  | **B2 Duration (seconds)** | 63 | **5.23** | **0.0005\*** |
| B3 | **B3 Peak frequency (kHz)** | 63 | **7.27** | **<0.0001\*** |
|  | **B3 Duration (seconds)** | 63 | **22.63** | **<0.0001\*** |
|  | Peak frequency pulses (kHz) | 63 | 1.32 | 0.266 |
|  | **Duration pulses (seconds)** | 63 | **13.80** | **<0.0001\*** |
|  | **Number of pulses (#)** | 63 | **11.21** | **<0.0001\*** |
|  | **Pulse rate (# pulses/duration pulses)** | 63 | **3.15** | **0.014** |
| **Phrase C** |  |  |  |  |
| C1 | **Peak frequency (kHz)** | 63 | **6.20** | **0.0001\*** |
|  | **Duration (seconds)** | 63 | **11.93** | **<0.0001\*** |
|  | Peak frequency lower tone (kHz) | 63 | 0.09 | 0.992 |
|  | **Duration lower tone (seconds)** | 63 | **2.93** | **0.020** |
|  | Peak frequency upper tone (kHz) | 61 | 0.17 | 0.974 |
|  | **Duration upper tone (seconds)** | 61 | **2.46** | **0.044** |

\*Significant at Bonferroni $\alpha = 0.0024$

##### i. Identification and Analysis of Syllables as Binary Variables

To define distinct syllables within and among songs we visually cataloged the syllables through inspection of the spectrograms with the bird IDs hidden (Searcy et al., 1985; Podos et al., 1992; González and Ornelas, 2009). This process was repeated three times by the same observer (ANB) for each song, resulting in 55 syllable types that were identified in all three iterations (Supplemental Material, Fig. S1). However, multiple syllables in this initial catalog were sung by only one bird, suggesting that we were over-splitting syllable types. We dealt with this in two ways. First, we removed syllables that were only sung by one individual; this reduced our number of syllable types to 42. Second, we binned syllables together that were produced in analogous syntax positions and had clear similarities in spectral shape, prominent frequency, harmonic structure, and duration (Supplemental Material, Fig. S1). To group these similar syllables together, we isolated syllables by splicing recordings into syllable units in Audacity 3.3.1 (https://audacityteam.org), such that each recording only consisted of one syllable and individual identity was hidden. To confirm that the visual binning process accurately captured acoustic similarity, we grouped syllables auditorily by listening to each representative syllable recordings at 0.25 speed and then at full speed. The visual and auditory groupings matched one another, resulting in 11 syllable groups that were scored as present or absent for each bird. Subsequent analyses were done on both this binned dataset and the full dataset with single incidence syllables removed (hereafter, full dataset).

To test for an effect of geographic distance on syllable use, we used the proportional abundance of each syllable by population to produce a Jaccard’s similarity matrix with the R package vegan (https://CRAN.R-project.org/package=vegan). Correlation between geographic distance and syllable use distance was tested with a Mantel test in XLSTAT (Lumivero, Burlington, MA, USA). To test for evidence of geographic structure in patterns of syllable use without a priori assignment of birds to populations, we converted individual syllable use to a binary matrix (1 = presence, 0 = absence) and ran a population structure analysis using the *snmf* function in the R package, LEA v3.11.6 (Frichot and François, 2015). Like the more computationally intensive programs STRUCTURE (Pritchard et al. 2000) and ADMIXTURE (Alexander et al., 2009), LEA is typically used in population genetics to estimate individual ancestry proportions from *K* populations or clusters, where the *K* is the number of populations that provides the best fit to the data. We follow González and Ornelas (2014) in extending this approach to the analysis of bird song. We ran ten replicates for each *K* value from 1 to 10, with a regularization parameter (*α*) value of 100 as recommended (Frichot et al. 2014). We chose the value of *K* with the lowest cross-entropy criterion value (Frichot et al. 2014) and evaluated the relationship between geographic origin and cluster membership.

##### ii. Quantitative Analysis of Syllables

We tested for an effect of population on each measure of duration and frequency with analysis of variance (ANOVA), with Bonferroni-corrected α = 0.0024. To better summarize and visualize spectral and temporal song differences among populations, we entered all measurements for phrases B-C into a principle component analysis (PCA). We excluded measurements for phrase A from this analysis because the high level of variation in the form and structure of syllables in phrase A reduced the explanatory power of the first two principle components by 47% relative to the analysis with phrases B and C only. We used population averages for the first three principial component (PC) scores to calculate Euclidian distance among populations, and tested for a correlation between geographic distance and song distance with a Mantel test. ANOVAs and PCA were run in JMP (SAS Institute Inc., Cary, NC, USA).

### Costa’s Hummingbird Song

Costa’s hummingbirds sing a one-phrase song consisting of a rapidly modulated tonal up and down sweep. We divided the songs into three parts: up-sweep, apex, and down-sweep (Fig. 1b). The up-sweep and down-sweep spanned an approximately 6 kHz range from 6 kHz to 12 kHz, with the apex at approximately 12 kHz. As for Anna’s, we chose one representative song per individual that had the highest signal to noise ratio and the lowest background noise. Due to the high frequency nature of the song, it degraded quickly and was affected by head movement. This made the beginning of the up-sweep and end of the down-sweep hard to define accurately. We therefore took a conservative approach and analyzed only the peak frequency and duration of the apex of the song. Whereas the duration of the whole song is variable within individual, the duration of the apex is stereotyped within individual (Williams and Houtman, 2008; ANB personal observation). Visual categorization of songs did not detect any discrete differences in signal form (Fig. 2b). Duration and frequency measurements were taken in Raven (Table 2) and we ran the same set of analyses described in Section ii. for quantitative measures of Anna’s song.

**Table 2.** Analysis of variance results for spectral and temporal measures of Costa’s hummingbird song.

|  | Measurement | *n* | *F* | *p* |
| --- | --- | --- | --- | --- |
| **Apex** | **Peak frequency (kHz)** | 45 | **2.95** | **0.032** |
|  | Duration (seconds) | 45 | 1.80 | 0.148 |
|  | High frequency (kHz) | 45 | 1.81 | 0.146 |
|  | **Low frequency (kHz)** | 45 | **2.75** | **0.041** |
|  | Delta frequency (kHz) | 45 | 1.27 | 0.297 |
| **Apex note 1** | **Peak frequency (kHz)** | 45 | **4.27** | **0.0057** |
|  | Duration (seconds) | 45 | 0.87 | 0.492 |
| **Apex note 2** | Peak frequency (kHz) | 45 | 1.28 | 0.295 |
|  | Duration (seconds) | 45 | 2.42 | 0.064 |

## RESULTS

### Evidence for Song Dialects in Anna’s

The Anna’s hummingbird populations sampled in this study varied in both syllable use, and in multiple measures of syllable frequency and duration. There was a significant positive association between syllable use distance and geographic distance for both the full (42 syllables; Mantel, *r =* 0.76, *r^2^* = 0.58, *P* < 0.0001; Fig. 3a) and the binned datasets (11 syllable groups; Mantel, *r =* 0.81, *r^2^* = 0.65; Supplementary Material, Fig. S2). We found a weak positive association between song distance and geographic distance for the spectral and temporal components of Anna’s song (Mantel test, *r* = 0.48, *r^2^* = 0.24, *P* = 0.076; Fig. 3b).

**Figure 3.**
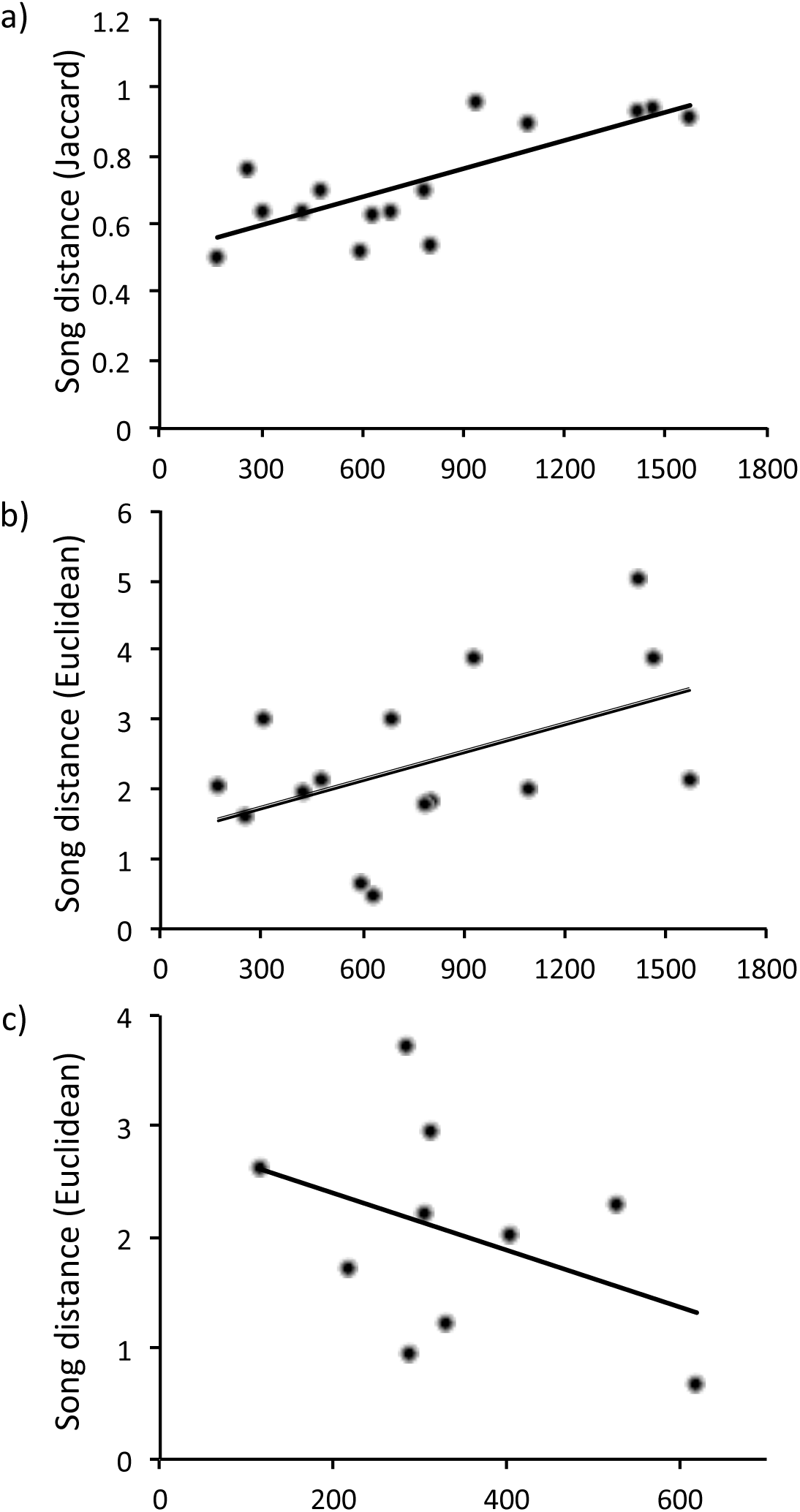
Effect of geographic distance on song distance in Anna’s (**a-b**) and Costa’s (**c**) hummingbirds. **a)** Geographic and song distance for syllable use in Anna’s (full dataset; Mantel, *r* = 0.76, *r^2^* = 0.58, *P* < 0.0001). **b)** Geographic and song distance for spectral and temporal components of Anna’s song (Mantel, *r* = 0.48, *r^2^* = 0.24, *P* = 0.076). **c)** Geographic and song distance for spectral and temporal components of Costa’s song (Mantel, *r* = 0.40, *r^2^* = 0.16, *P* = 0.26). Geographic distance estimated as linear distance in kilometers from center of each sampling locality; song distance calculated as Jaccard distance for syllable use and Euclidian distance for spectral and temporal measures.

Patterns of syllable use were spatially structured. For both full and binned datasets, *K*=4 clusters had the lowest cross-entropy criterion value, with similar proportional assignments of individual birds to clusters (Fig. 4; Supplementary Material, Fig. S3). In the full dataset, 12 of 13 Seattle (SE) birds were assigned to a Seattle-limited cluster (Cluster 1) and all Mendocino (ME) and Santa Ynez (SY) birds were assigned to a single cluster (Cluster 3; Fig.4). Henderson (HE), San Francisco (SF), and Riverside (RI) birds were less differentiated in syllable use; individuals from all three populations, together with one bird from SE, were represented in Cluster 2. Riverside was distinct from all other populations in lacking representation in Cluster 3, and Cluster 4 was unique to RI and SF (Fig. 4).

**Figure 4.**
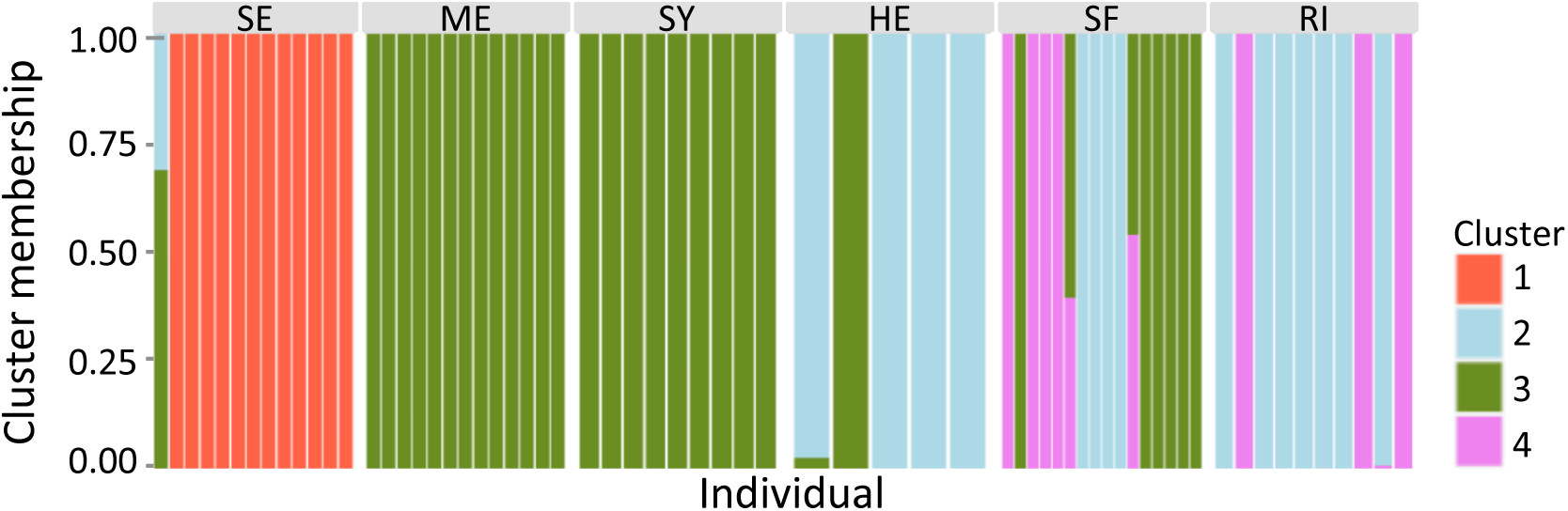
Geographically structured patterns of syllable use in Anna’s hummingbird based on full dataset. Bars are individual birds, colors indicate proportional assignment to *K* = 4 clusters. SE, Seattle; ME, Mendocino; SY, Santa Ynez; HE, Henderson; SF, San Francisco; RI, Riverside.

There was a significant effect of population on 13 of the 21 spectral and temporal measures of Anna’s song; seven of these survived Bonferroni correction (α = 0.0024; Table 1; Fig. 5; Supplementary Material, Fig. S4). We ran post hoc Tukey HSD tests on these seven variables to determine which populations were driving each result (Fig. 5). Although the number of significant pairwise population contrasts differed between variables, some general patterns were evident. As for the syllable use clustering analysis, SE birds were most consistently differentiated for significant temporal measures (shorter duration for Phrase B elements B2 and B3 with fewer B3 pulses; longer duration for Phrase C element C1; Fig. 5a-e), and ME and SY were consistently similar (Fig. 5a-g). Population differences were more variable for significant frequency measures, with higher B3 frequency in HE, RI and SE relative to ME and SY (Fig. 5g), and higher C1 frequency in RI only (Fig. 5f).

**Figure 5.**
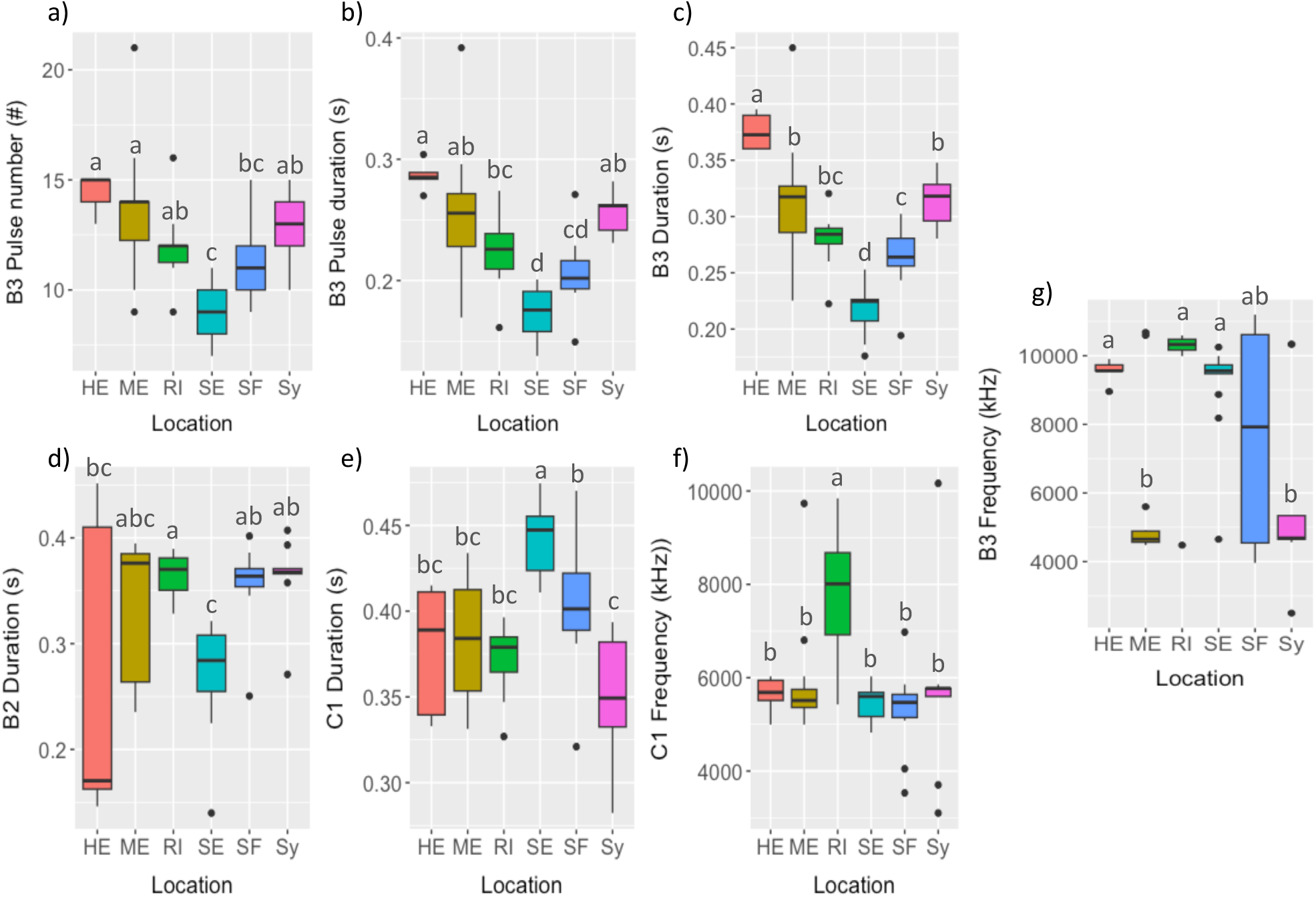
Geographic variation in spectral and temporal measures of Anna’s hummingbird song. Populations that do not share letters are significantly different (Tukey HSD test). All ANOVA p-values ≤ 0.0005 (see Table 1). HE, Henderson; ME, Mendocino; RI, Riverside; SE, Seattle; SF, San Francisco; SY, Santa Ynez.

Principle component analysis of all Phrase B and C variables similarly separated SE from the other five populations. The first two components explained 45% of the total variance with duration and frequency positively loaded on PC1 and PC2, respectively (Fig. 6a). Mean scores for SE were negative on both components, reflecting shorter duration and lower frequency relative to all other populations. PC1 also distinguished SF and RI from SY and HE, with more positive mean scores indicative of longer durations in the latter two populations (Fig. 6a).

**Figure 6.**
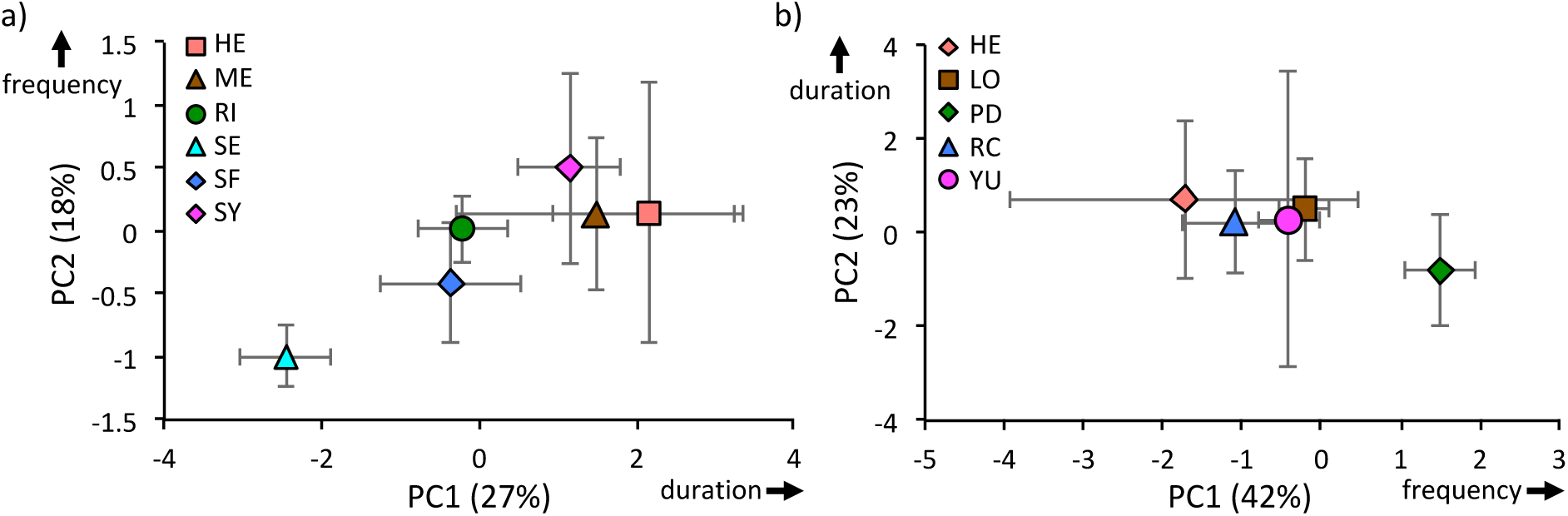
Principle component analysis of spectral and temporal measures of Anna’s and Costa’s hummingbird song. Plots of population mean scores for first (PC1) and second (PC2) principle components from, **a)** six Anna’s populations with duration positively loaded on PC1 and frequency positively loaded on PC2 and **b)** five Costa’s populations with frequency positively loaded on PC1 and duration positively loaded on PC2. Axis labels include percent variance explained by each PC; error bars are standard deviations. HE, Henderson; ME, Mendocino; RI, Riverside; SE, Seattle; SF, San Francisco; SY, Santa Ynez; LO, Lompoc; PD, Palm Desert; RC, Rancho Cucamonga; YU, Yuma.

### Little Geographic Variation in Costa’s Song

There was no significant association between song distance and geographic distance among Costa’s populations (Mantel test, *r* = 0.40, *r^2^* = 0.16, *P* = 0.26). Indeed, more distant populations tended to have lower song distances (Fig. 3c). Song form was invariant based on visual evaluation of spectrograms (see Fig. 2b).

There was a significant effect of population on three of the nine spectral and temporal measures of Costa’s song, but none survived Bonferroni correction (α = 0.0056; Table 2; Fig. 7). Apex frequency measures tended to be higher in Palm Desert (PD) and lower in Henderson (HE) (Fig. 7), whereas all duration measures were more variable within than between populations (Supplementary Material, Fig. S5). Principle component analysis of all nine variables recovered similar patterns (Fig. 6b). The first two components explained 65% of the total variance. Frequency was positively loaded on PC1, which partially separated PD from other populations. All populations clustered together on PC2, on which duration was positively loaded (Fig. 6b).

**Figure 7.**
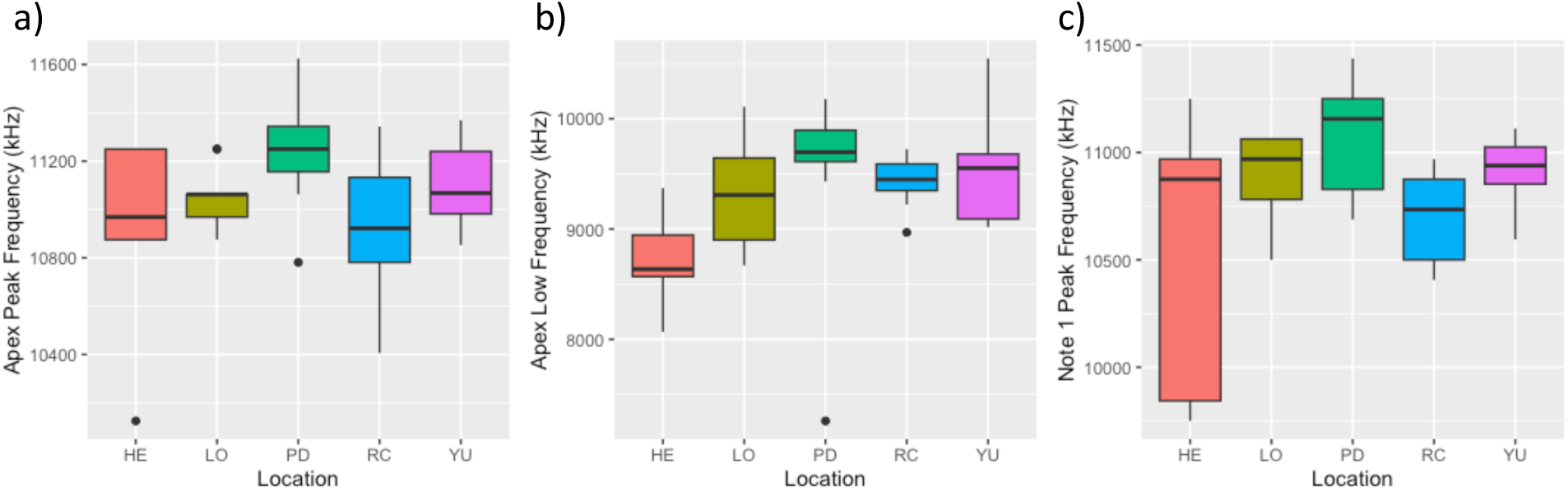
Geographic variation in spectral measures of Costa’s hummingbird song. All ANOVA p-values > 0.0056 (see Table 2). HE, Henderson; LO, Lompoc; PD, Palm Desert; RC, Rancho Cucamonga; YU, Yuma.

## DISCUSSION

We tested for the presence of song dialects in a pair of closely related hummingbirds that differ in song complexity. Consistent with our expectation that a more complex song should provide more opportunities for copying errors and the accumulation of population variants, we found robust evidence for song dialects in Anna’s and contrastingly little in Costa’s. In Anna’s, we recovered a strong positive association between geographic distance and song distance based on syllable use, and a weaker but still positive association for song distance based on spectral and temporal measures. This pattern of isolation-by-distance is consistent with the notion that dialect formation is a byproduct of cultural drift – the stochastic accumulation of acoustic differences between populations, akin to differentiation in allele frequencies due to genetic drift.

In Anna’s, the song of birds from Seattle was differentiated from all other populations in terms of both syllable use, and quantitative measures of song. In particular, the absence of a B2 syllable and the presence of a unique C1 syllable were the clearest differences between Seattle and the other populations we sampled. Seattle is the northernmost population we recorded (Fig. 2a), and is part of a northward range expansion in Anna’s hummingbirds that has occurred in the last two decades. This recent range expansion is thought to be facilitated by human activities that provide additional food sources for hummingbirds. These include, extensive use of hummingbird feeders, plantings that attract hummingbirds, and an increase of non-native flowers such as tree tobacco (*Nicotiana glauca*). In addition, higher overwinter survival is associated with urbanization (Zimmerman, 1973; Grieg, 2017). We speculate that syllable use differences in the Seattle population, particularly the apparent loss of syllable B2, are due to a founder effect on song. We note, however, that the largest geographic distances in our sampling scheme were between Seattle and other populations. Therefore, some of the quantitative differences in spectral and temporal measures of Seattle birds’ song may be points on a continuum of variation along the West Coast of North America. A complete understanding of the effect of northward range expansion on Anna’s song awaits sampling of intervening populations in Oregon and Southern Washington State.

The close association in syllable use between Mendocino in Northern California and Santa Ynez in Southern California could not be explained by geographic proximity. The annual movement patterns of Anna’s hummingbirds are not well defined so designating how much and historically when populations interacted with one another is unclear perhaps leading to SY and ME populations to group together. Adult males hold display territories from January through June, but then leave and where they go is unknown. In contrast, juveniles tend to stay in the breeding areas June-August. Since males leave in June and females rarely sing, it is unclear who the later-hatched juveniles are learning their songs from. Even earlier in the breeding season, it seems that nests are not placed on or near male display territories (ANB personal observation). Moreover, there is evidence for expansion and subsequent contraction of the eastern range of Anna’s (Zimmerman, 1973). The effect of seasonal and longer-term movements on song learning and spatial distribution of song variants clearly requires further investigation in Anna’s.

The relatively less complex structure of Costa’s song, including the absence of discreet syllables, meant that our search for among-population differences in song was based on far fewer variables than in Anna’s. We found no qualitative evidence for variation in song structure and there was no relationship between geographic and song distance based on quantitative measures. Given that the upper range of straight line distances between Costa’s populations was less than half of that for Anna’s populations, it is possible that an effect of geography on song might emerge with additional sampling. However, the tendency for song distance to decrease with spatial distance suggests the addition of more distant populations would not uncover a pattern of isolation-by-distance in Costa’s song.

Costa’s songs from Palm Desert tended to differ from other populations in having a higher frequency range. Notably, Palm Desert songs were recorded from directed short range song produced during dynamic, undulating flight (shuttle displays), whereas recordings from other populations were from males singing while perched. Given the absence of frequency differences in other Costa’s populations, we think it likely that we uncovered a previously undocumented effect of context on male song rather than a consistent population difference.

In summary, we conducted the first macrogeographic study of geographic variation in the songs of two closely related species of hummingbirds, a song-learning lineage that is generally underrepresented in studies of song dialects. We found extensive geographic variation in Anna’s song and contrastingly little in Costa’s. A critical next step in this line of research is to determine whether the statistically significant population differences in Anna’s song are biologically significant using field playbacks of population-matched vs. –mismatched songs. Conversely, the absence of statistical significance does not guarantee that Costa’s songs from different populations are equivalent from the birds’ perspectives. Therefore, playback experiments are similarly well motivated in Costa’s.

## ACKNOWLEDGEMENTS

This research was supported by grants from the Animal Behavior Society and Western Field Ornithologists to ANB, and by equipment from the National Academy of Sciences-Keck Foundation Initiative’s Project ECAT. We thank David Rankin, Ilana Kornfeld, and Shaina Hammer for field support.

## Notes

### Competing Interest Statement

The authors have declared no competing interest.

