## Supplemental Figures S1-5 for "Geographic variation in the songs of two closely related song-learning species, Anna’s and Costa’s hummingbird (*Calypte anna, C. costae*)"

### SUPPLEMENTAL MATERIAL

**Figure S1**

**Phrase 1**

A group

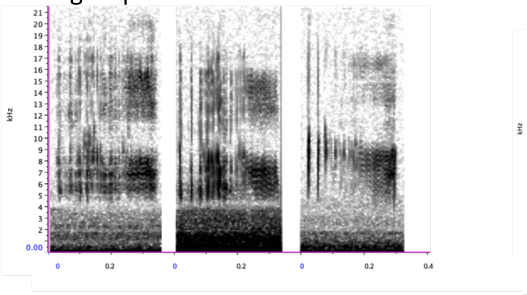

B group

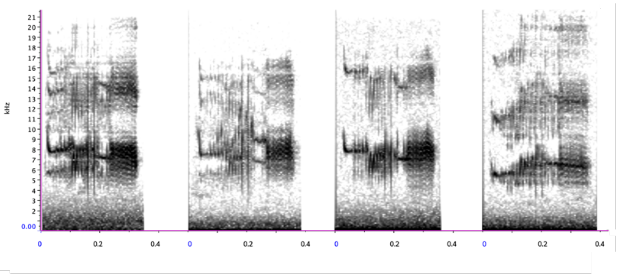

C group

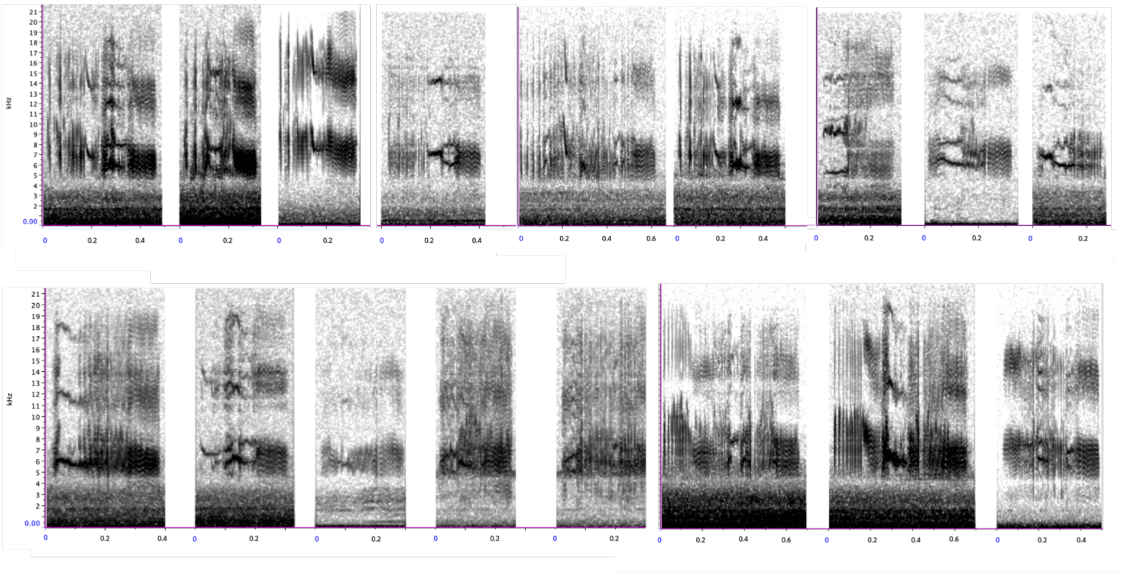

D group

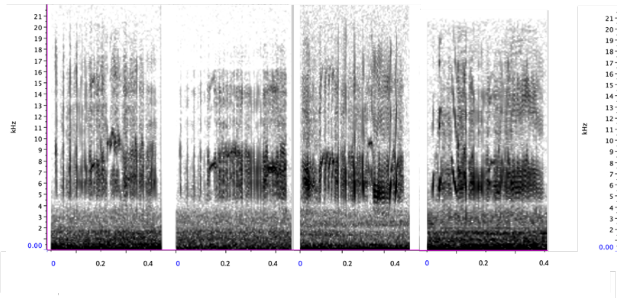

F group

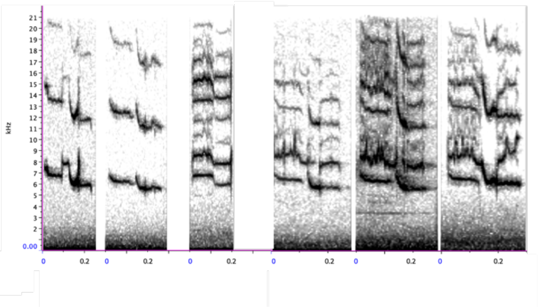

### Figure S1 (continued)

**Phase 2**

B1 group

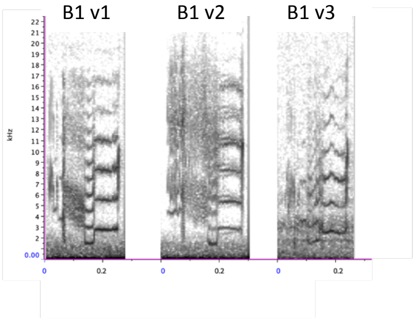


B2a group

B2b group

B2c group

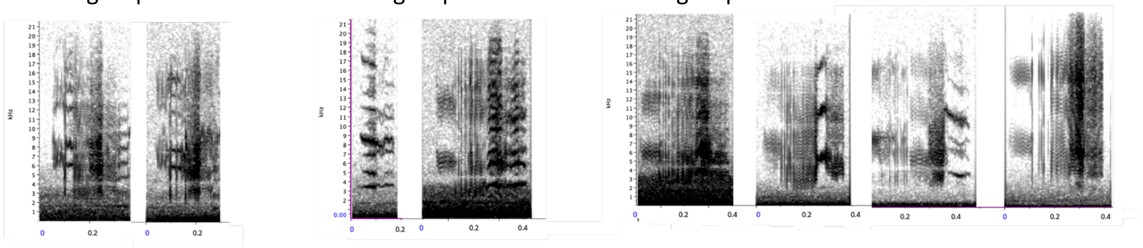

B3 group

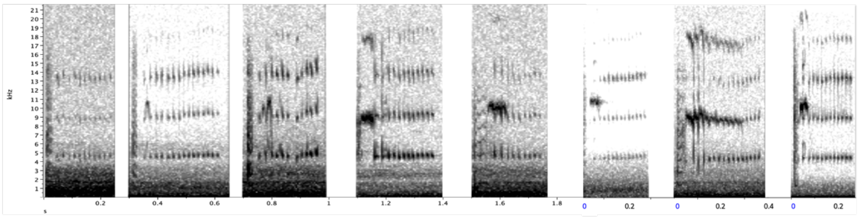

**Phase 3**

C1 group

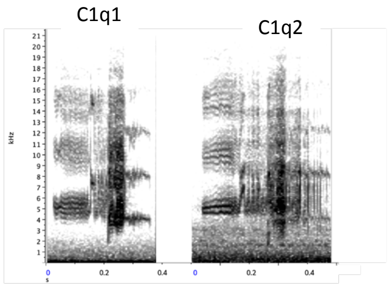


**Figure S1.** Syllable groupings in Anna's hummingbird song. Fifty-five syllables were identified and were binned into 11 groups.

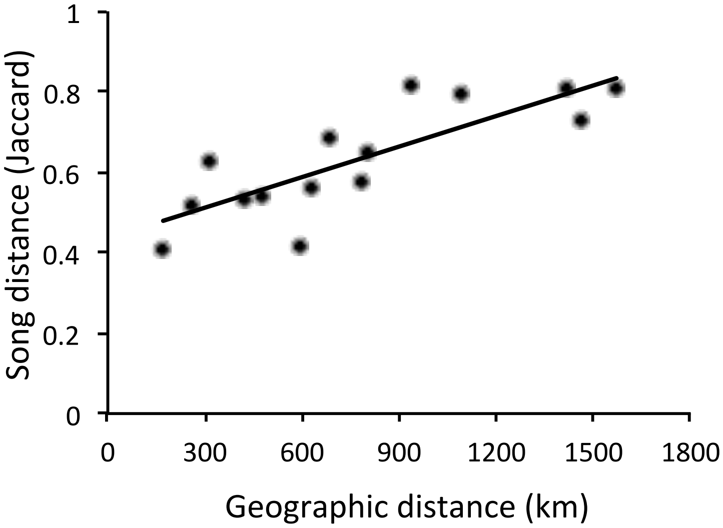


**Figure S2.** Effect of geographic distance on song distance for syllable use in Anna's hummingbirds (binned dataset; Mantel, $r = 0.81$, $r^2 = 0.65$, $P < 0.0001$). Geographic distance estimated as linear distance in kilometers from center of each sampling locality; song distance calculated as Jaccard distance.

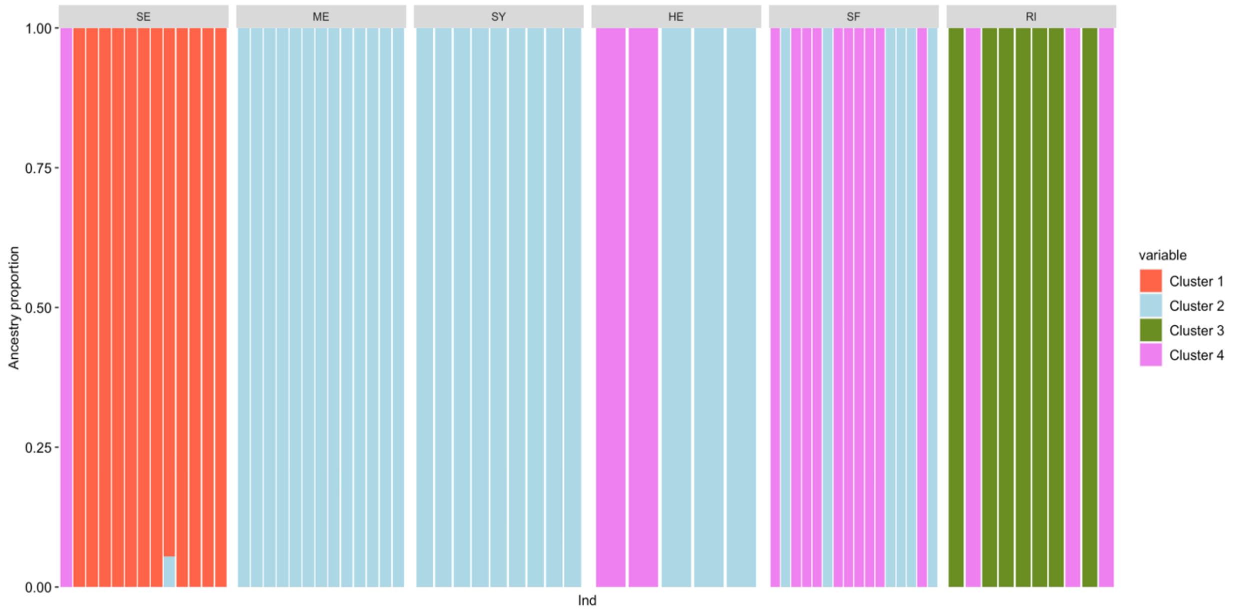


**Figure S3.** Geographically structured patterns of syllable use in Anna's hummingbird based on binned data. Bars are individual birds, colors indicate proportional assignment to $K = 4$ clusters. SE, Seattle; ME, Mendocino; SY, Santa Ynez; HE, Henderson; SF, San Francisco; RI, Riverside.

**Figure S4**

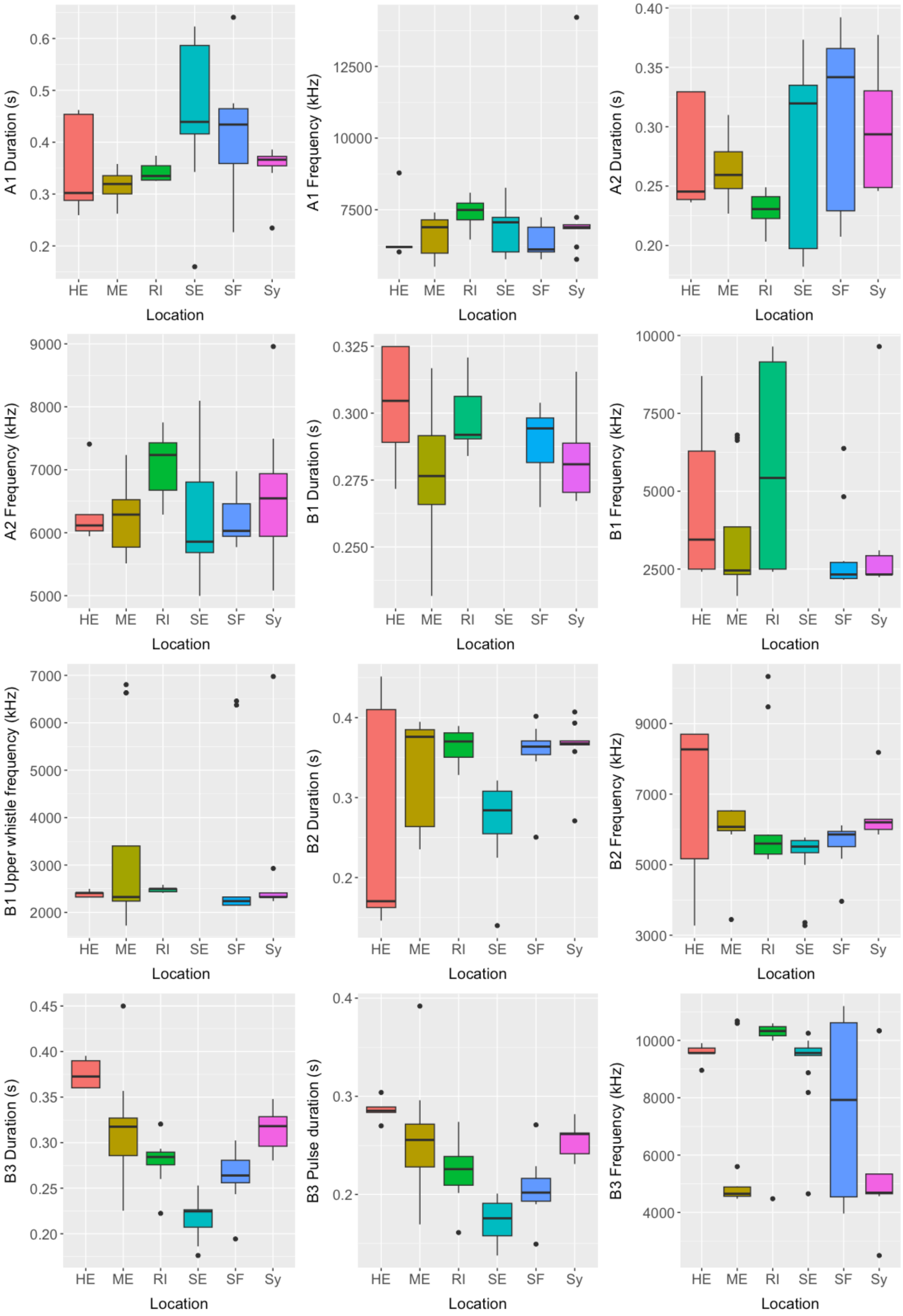
A1 Duration (s)
A1 Frequency (kHz)
A2 Duration (s)
A2 Frequency (kHz)
B1 Duration (s)
B1 Frequency (kHz)
B1 Upper whistle frequency (kHz)
B2 Duration (s)
B2 Frequency (kHz)
B3 Duration (s)
B3 Pulse duration (s)
B3 Frequency (kHz)
HE
ME
RI
SE
SF
Sy
Location

**Figure S4 (continued)**

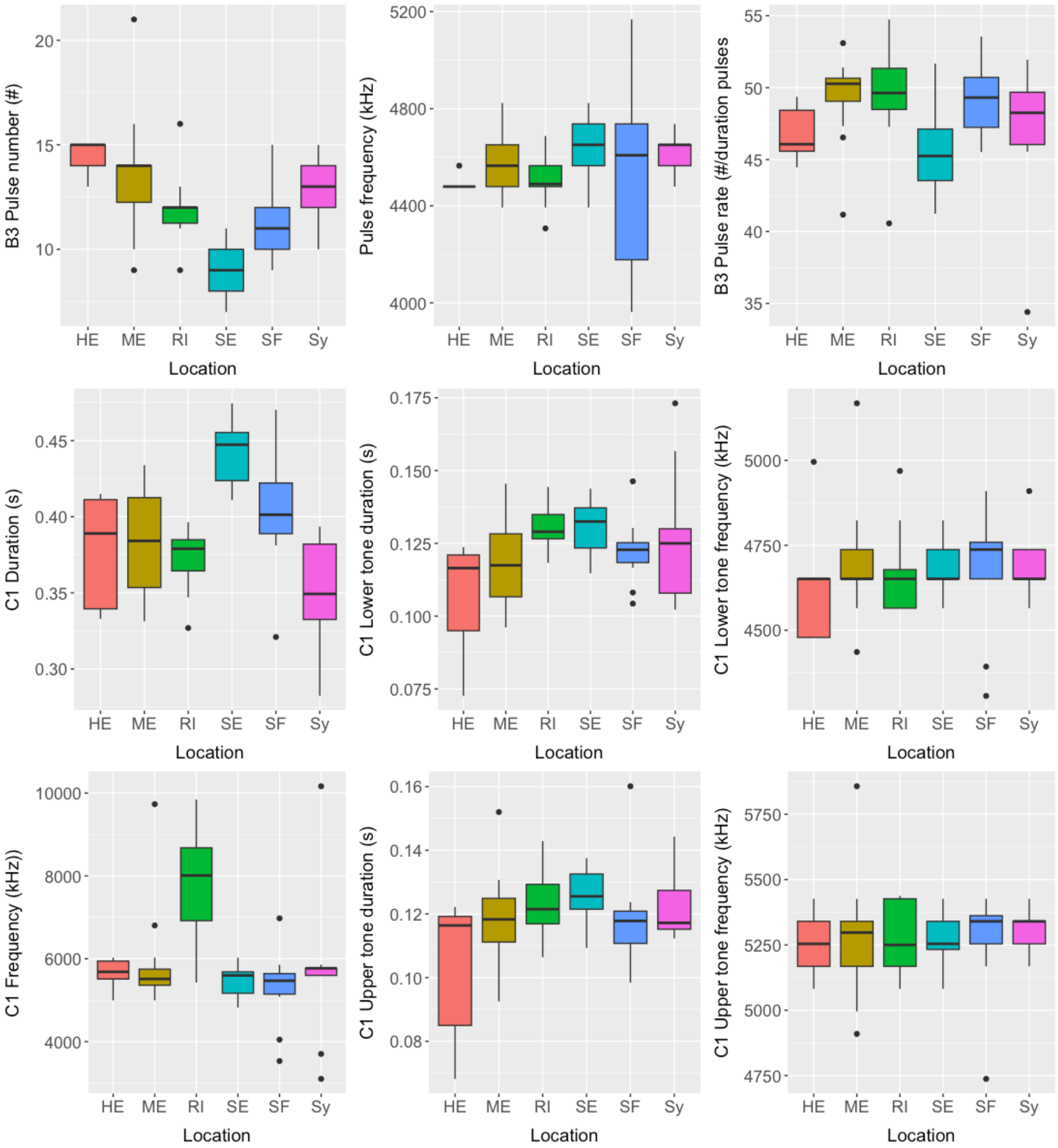


**Figure S4.** Twenty-one spectral and temporal measures of Anna's hummingbird song from six populations. HE, Henderson; ME, Mendocino; RI, Riverside; SE, Seattle; SF, San Francisco; SY, Santa Ynez.

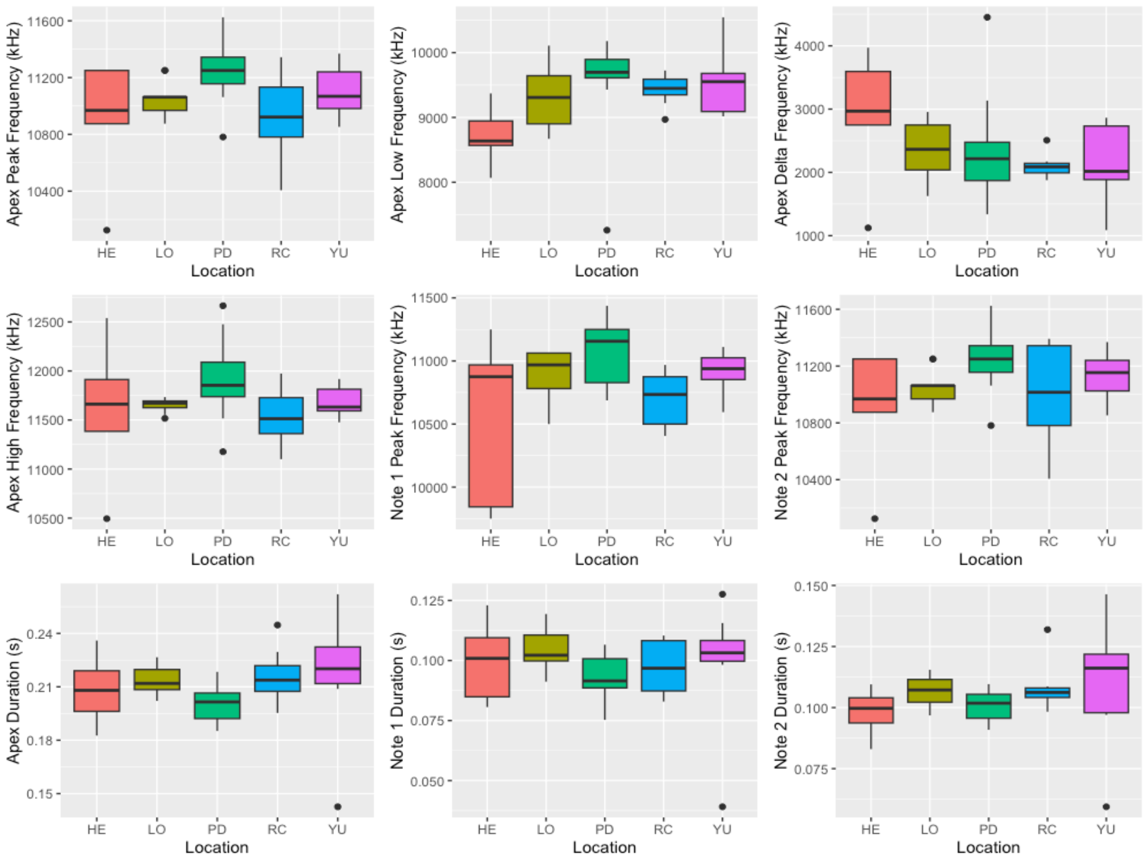


**Figure S5.** Nine spectral and temporal measures of Costa's hummingbird song from five populations. HE, Henderson; LO, Lompoc; PD, Palm Desert; RC, Rancho Cucamonga; YU, Yuma.
